# Genome Report: Genome of the Amazon Guppy (*Poecilia bifurca*) reveals conservation of sex chromosomes and dosage compensation

**DOI:** 10.1101/2025.05.19.654947

**Authors:** Lydia J.M. Fong, Bernadette D. Johnson, Iulia Darolti, Benjamin A. Sandkam, Judith E. Mank

## Abstract

The Amazon guppy, *Poecilia bifurca*, is a small live-bearing fish. Closely related species, such as *P. reticulata, P. picta*, and *P. parae* all share the same sex chromosome system, but with substantial diversity in the degree of Y degeneration and the extent of X chromosome dosage compensation. We built a female (XX) draft genome with 55X coverage of PacBio HiFi data, resulting in a 785 Mb assembly with 94.4% BUSCO completeness. Using this genome, we found that *P. bifurca* shares the same sex chromosomes as related species and shows substantial Y chromosome degeneration. We combined this with RNA-Seq data and find similar expression of X-linked genes between sexes, revealing that *P. bifurca* also exhibits complete X chromosome dosage compensation. We further identify 11 putative autosome-to-Y gene duplications, five of which show gene expression in guppy male germ cells.

## Introduction

Fishes in the genus *Poecilia* have been extensively studied in the context of sexual selection and sexual conflict (Endler 1980), life-history evolution (Meredith et al. 2011), and are an emerging model system for sex chromosome evolution. Several closely related species in the genus share the same sex chromosome system but with extensive differences in the extent of Y chromosome degeneration and X chromosome dosage compensation. *Poecilia reticulata* has homomorphic sex chromosomes with little degeneration of the Y chromosome in the region homologous to the X (Darolti et al. 2019; Du et al. 2025; Wright et al. 2017), while the non-recombining region of the Y chromosome in its sister species, *P. wingei*, has expanded substantially and is the largest chromosome in the genome (Nanda et al. 2014). *P. parae* and *P. picta* belong to a sister clade yet have heteromorphic sex chromosomes, extensive Y degeneration, and complete X chromosome compensation (Darolti et al. 2019; Fong et al. 2023; Metzger et al. 2021; Sandkam et al. 2021). Despite the difference in the state of Y chromosome degeneration and dosage compensation, these four species share a recent, single origin of their sex chromosome (Fong et al. 2023). This begs the question of why there is so much variation between species and what might we expect to observe in other members of this genus.

The Amazon guppy (*Poecilia bifurca*) is sexually dimorphic (Figure 1a) and is in the same clade as *P. picta* and *P. parae* (Figure 1b; Meredith et al. 2011; Pollux et al. 2014; Rabosky et al. 2013; Rabosky et al. 2018) but its ecology, including life history and mating behaviour, are less studied compared to its close relatives (Breden et al. 1999; Rosen and Bailey 1963). Furthermore, it lacks a reference genome, and any potential sex chromosomes have not been identified. We might predict that *P. bifurca* possesses a degenerated Y chromosome shared with *P. picta* and *P. parae*. However, frequent sex chromosome turnover has been observed among closely related fish species (Vicoso 2019), for example in sticklebacks (Ross et al. 2009), Lake Tanganyika cichlids (Gammerdinger & Kocher 2018; Roberts et al. 2009), and medaka (Myosho et al. 2015), suggesting that this is also a possibility for *P. bifurca*.

**Figure 1.**
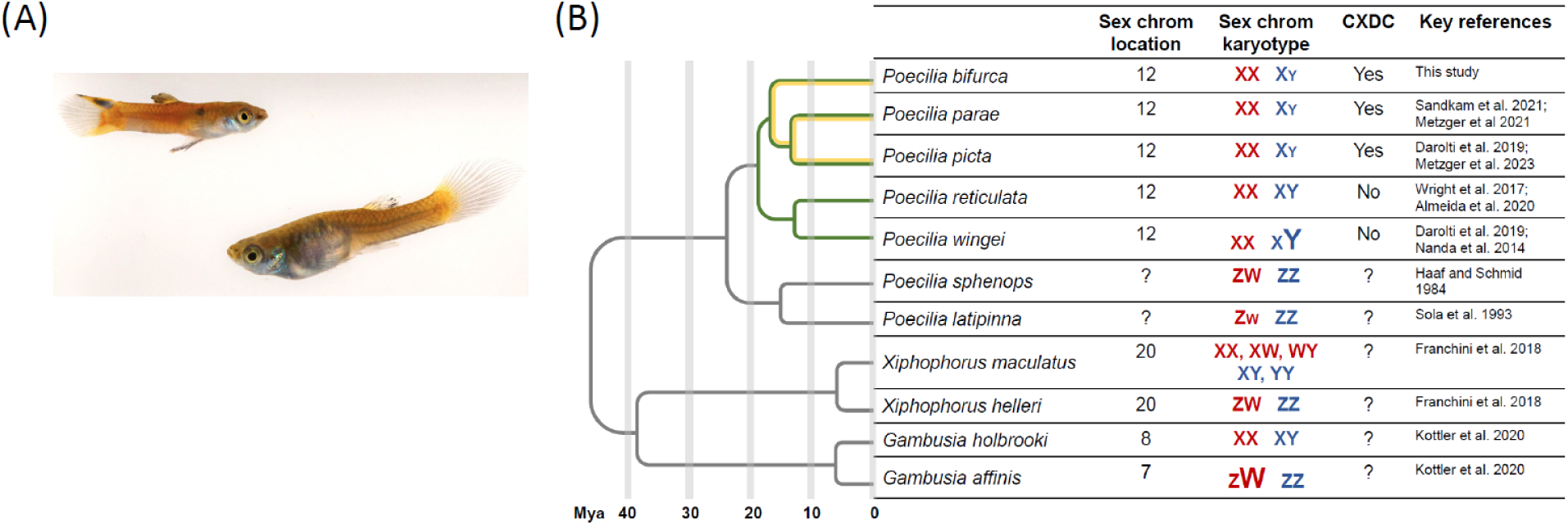
Photo of adult *P. bifurca* and the phylogeny of closely related species. A) A male individual is depicted on the left, a female on the right. B) Phylogeny from Rabosky et al. (2018) and Rabosky et al. (2013). Maximum likelihood origin of the sex chromosome system in *P. bifurca* and its close relatives is shown in green, and complete X chromosome dosage compensation (CXDC) shown in yellow. Included is the sex chromosome location on guppy chromosome number, relative size differences of the sex chromosomes with female karyotypes in red and male in blue, evidence of complete X chromosome dosage compensation, and references.

To address this question, we first built a draft female genome for *P. bifurca* using PacBio long-read sequencing of an individual female (XX). Using short read DNA data from males and females, we determined that *P. bifurca* shares the same sex chromosome with its close relatives (guppy chromosome 12), and that the Y chromosome is highly degenerated, similar to *P. picta* and *P. parae*. Using expression data from males and females, we show that *P. bifurca* also shares complete X chromosome dosage compensation with *P. picta* and *P. parae*. Finally, we use coverage estimates for both sexes and polymorphism data to identify 11 putative autosome-to-Y chromosome duplications and explore the expression of these loci.

## Materials and Methods

### Sample collection and sequencing methods

All individuals were kept in temperature-controlled chambers at 26-28°C, and tanks were kept at a pH range from 6.0-7.0. Individuals for sequencing were flash frozen in liquid nitrogen and stored in -80°C until sample extraction. For genome assembly, high molecular weight (HMW) DNA was collected from one female (25.1 ng/μL) using the New England Biotechnological Lab Monarch Kit for PacBio Sequel II SMRT 8M. HMW DNA fragments longer than 20kb were used for CCS HiFi library prep, and sequencing was done using one SMRT cell to the resulting average genomic coverage was 55X.

For short-read DNA sequencing, we extracted DNA from three males and three females using the Qiagen DNeasy Blood and Tissue Kit (31.5 – 187.7 μg/μL). Libraries were prepared using a PCR-free method and sequenced on a single lane of a NovaSeq S4 PE150 flow cell, generating an average coverage of ∼62.3X (Supplemental Table 1). For RNA-sequencing, we collected somatic (tail and head) tissue from three adult males and three adult females, and used the Qiagen RNeasy Kit to extract total mRNA (RIN ≥ 8). Libraries were prepared using the NEB Ultra II Directional poly(A) mRNA kit and sequenced on a single lane of a Novaseq SP flowcell for 150bp paired-end reads (PE2x150bp). All library preparations and sequencing were done at Toronto SickKids Hospital Sequencing Centre.

For both the DNA and RNA short read data, we followed the quality assessment protocol from Darolti et al. (2019). Briefly, sample quality for Illumina sequences was assessed with FastQC v.10.1 (Andrews 2010; http://www.bioinformatics.babraham.ac.uk/projects/fastqc/, last accessed on July 8, 2023). Adaptors were removed and trimmed using Trimmomatic v.0.36, using the universal adapter for RNA sequences with the commands LEADING: 3 TRAILING: 3 SLIDINGWINDOW:4:15 MINLEN:50, PHRED 33. DNA sequences were trimmed using TruSeq3-PE-2 adapter following the same commands.

### Genome assembly and repeat annotation

The PacBio sequences were corrected (i.e. consensus of overlapping reads), trimmed, and assembled using CANU v.2.2 (Koren et al. 2017) using a maximum error allotment of 4.5%, an estimated genome size of 750 Mb, and all other parameters set to default. We then used Pilon v.1.22 (Walker et al. 2014) with *--fix all* and *--mindepth 0.5* to correct the long-read CANU assembly with the female *P. bifurca* Illumina DNA sequences. Finally, the draft genome assembly was oriented to the female *P. picta* reference genome (Metzger et al. 2023) using RagTag v.2.1.0 (Alonge et al. 2022) following the *ragtag_scaffold.py* script. We then ran BUSCO v.5.3.2 with the cyprinodontiformes_odb10 database (Simão et al. 2015) to check for genome assembly completeness (∼94.4%; Table 1). Lastly, we followed the Earl Grey (Baril et al. 2024) pipeline with default parameters to identify repeat content.

**Table 1.**
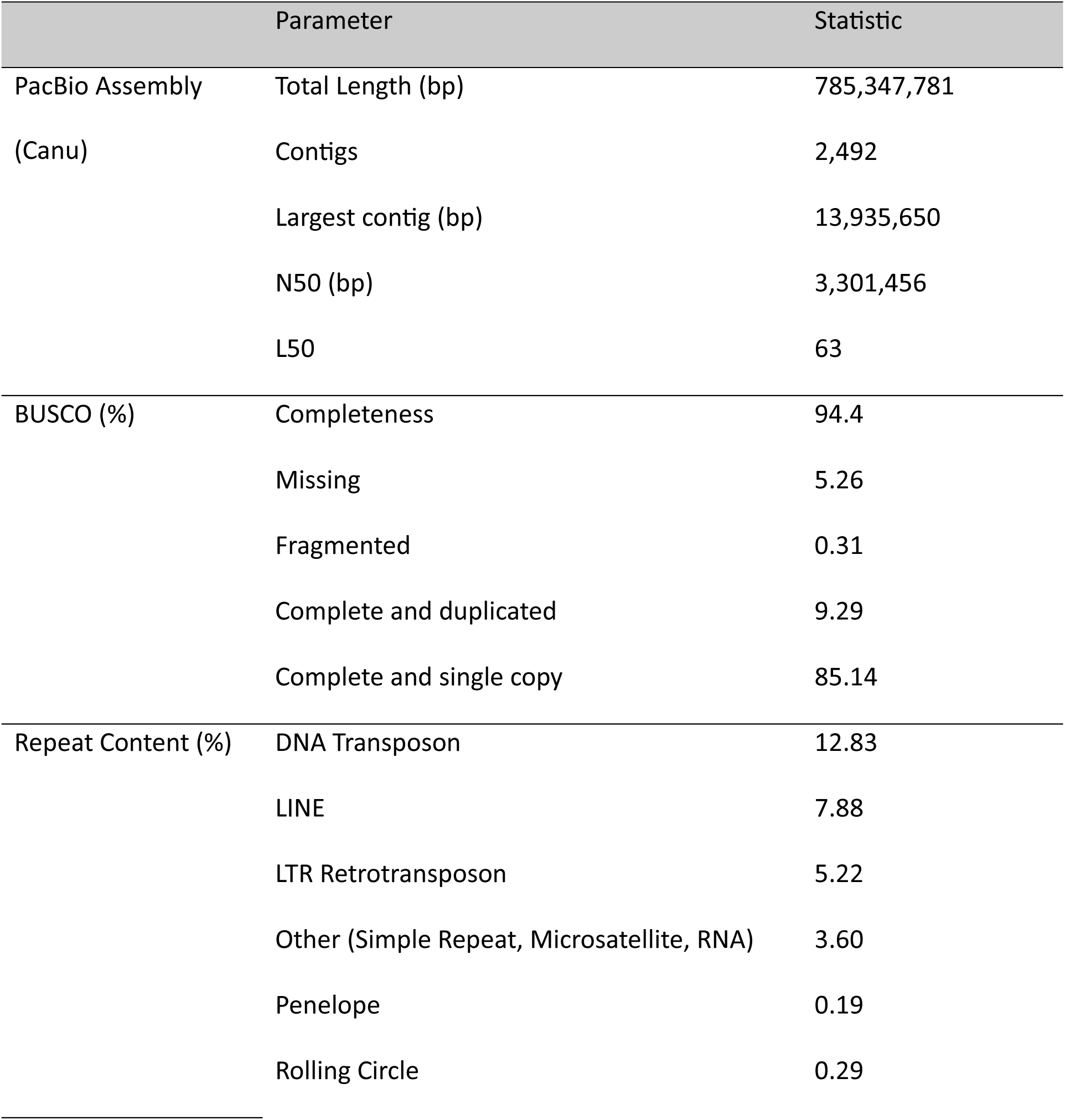
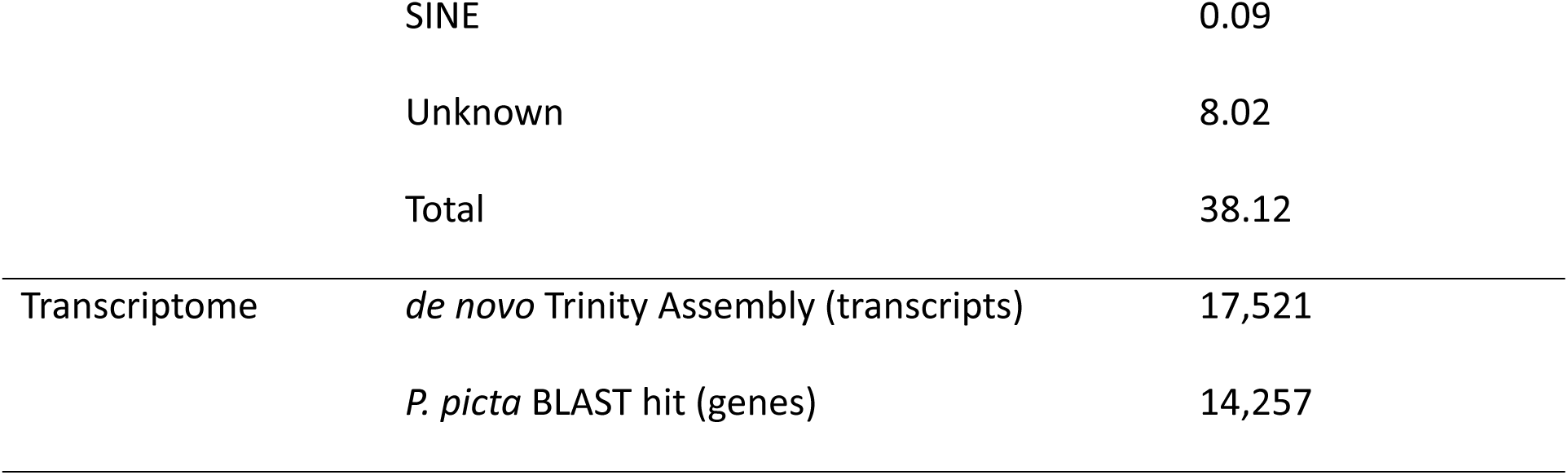
Assembly statistics for the draft genome, its repeat content, and for the transcriptome. BUSCO results using the cyprinodontiformes_odb10 database are reported as percentages. Numbers reported in the repeat content represents percentage of the genome that is composed of the classified repeat DNA. The number of *de novo* transcripts assembled from Trinity and the number of transcripts that BLAST to the *P. picta* genome annotation are reported.

|  | Parameter | Statistic |
| --- | --- | --- |
| PacBio Assembly<br>(Canu) | Total Length (bp) | 785,347,781 |
|  | Contigs | 2,492 |
|  | Largest contig (bp) | 13,935,650 |
|  | N50 (bp) | 3,301,456 |
|  | L50 | 63 |
| BUSCO (%) | Completeness | 94.4 |
|  | Missing | 5.26 |
|  | Fragmented | 0.31 |
|  | Complete and duplicated | 9.29 |
|  | Complete and single copy | 85.14 |
| Repeat Content (%) | DNA Transposon | 12.83 |
|  | LINE | 7.88 |
|  | LTR Retrotransposon | 5.22 |
|  | Other (Simple Repeat, Microsatellite, RNA) | 3.60 |
|  | Penelope | 0.19 |
|  | Rolling Circle | 0.29 |

|  |  |
| --- | --- |
| SINE | 0.09 |
| Unknown | 8.02 |
| Total | 38.12 |
| Transcriptome | *de novo* Trinity Assembly (transcripts) | 17,521 |
|  | *P. picta* BLAST hit (genes) | 14,257 |

### Transcriptome assembly

We first used HISAT2 v.2.1.0 (Kim et al. 2019) to align the RNA sequences to our draft *P. bifurca* genome and then used Trinity v.2.11.0 (Grabherr et al. 2011) with default parameters to build our *de novo* transcriptome. Reads were combined from all samples, then we filtered the transcriptome by removing unpaired and discordant alignments using the built-in Trinity script *align_and_estimate_abundance.pl* that maps the *de novo* transcriptome using Bowtie2 (Langmead et al. 2009). We filtered the transcriptome noncoding RNA by removing transcripts with BLAST hits to the *Oryzias latipes* ncRNA database (Ensembl: ASM223467v1; Hunt et al. 2018). The transcriptome was further filtered to remove coding regions within transcripts without an open reading frame and those with open-reading frames smaller than 150 bp using Transdecoder v.5.5.0 (Haas et al. 2013; http://transdecoder.github.io, last accessed November 24, 2023) with default parameters. The Transdecoder contigs were further assembled with CAP3 (Huang & Madan 1999) to build the final transcriptome. Genes from the transcriptome were annotated using BLAST against the *P. picta* female genome annotation and using the top blast hit.

### Identifying the sex chromosomes and dosage compensation

Degeneration of the Y results in reduced read coverage of males compared to females for the X chromosome. To determine the location of the sex chromosomes and the extent of Y degeneration, we used the male and female Illumina DNA data to identify the sex chromosomes. First, we mapped short reads to the *P. bifurca* draft genome assembly using bwa *aln* and *samse* v0.6.1 (Li & Durbin 2009), then found unique mapping sites using grep ‘XT:A:U’. We then sorted the aligned reads using SAMTools v.1.3.1 (Li et al. 2009) and extracted the coverage for each individual using soap.coverage v.2.7.7 (https://bio.tools/soap; Li et al. 2008). We calculated the average coverage for males and females separately and then compared male to female (M:F) coverage (log_2_ average male coverage/log_2_ average female coverage). We also used Bowtie2 v.1.1.2 (Langmead et al. 2009) to map the reads of all samples to the draft genome assembly. We re-analyzed *P. picta* and *P. parae* using the same pipeline to compare sex chromosome coverage across the species. These samples included three female and three male *P. picta* individuals from Darolti et al. (2019) and three females and three males of the parae morph of *P. parae* from Sandkam et al. (2021). Finally, we used the HAWK v. 1.7.0 pipeline (Rahman et al. 2018) to count and identify male-specific k-mers (Y-mers) for each species using counts from the Bonferroni-corrected file. We identified shared Y-mers with normalized coverage greater than 20X coverage (the threshold used to detect excess unique Y-mers relative to female-specific k-mers) across *P. bifurca*, *P. picta*, and *P. parae* to test for shared ancestry of the Y chromosome.

To determine the extent of X chromosome dosage compensation, we followed the approach of Darolti et al. (2019) to calculate allele-specific expression (ASE). First, we aligned the RNA-seq data to the *P. picta* female reference genome using STAR align v. 2.7.11 (Dobin et al. 2013) with default parameters and *–outFilterMultimapNmax 1*. We then called single-nucleotide polymorphisms (SNPs) for males and females separately using SAMTools v.1.3.1 (Li et al. 2009) *mpileup* and filtered the vcf file using VarScan v2.3.9 (Koboldt et al. 2009) *--min-coverage 2 -- min-ave-qual 20 --min-freq-for-hom 0.90 --p-value 1 --strand-filter 0 --min-var-free 1e-10*. We removed clusters of more than 5 SNPs in 100 bp windows to avoid bias in our ASE estimations from the preferential assignment of reads to the reference allele and removed triallelic SNPs.

We expect most genes to have biallelic expression (equal expression of the alleles from both chromosomes) and thus expect ∼0.5 probability to recover reads from either chromosome. Therefore, we tested for ASE by identifying significant deviations from 0.5 using a two-tailed binomial test (*p* < 0.05) on the final filtered SNP dataset. We called SNPs as ASE if a minimum of 70% of the reads came from one chromosome and kept identified genes with ASE that had at least one SNP with a consistent ASE pattern across all heterozygous samples. We then used a Chi-squared test examine differences in ASE patterns between the autosomes and the sex chromosomes in males and females. Finally, we used HTSeq v.2.0.5 (Putri et al. 2022) to count reads for the transcripts assembled from our *de novo* transcriptome and normalized the counts using gene length to get gene expression and compared M:F expression.

### Y gene duplication analysis

We followed the pipeline from Lin et al. (2022) to identify putative autosome-to-Y chromosome gene duplications using a combination of M:F F_ST_, M:F coverage, and M:F SNP density. Once duplicated from an autosome to the Y, the accumulation of Y-specific SNPs will produce elevated M:F F_ST_ and M:F SNP density > 1 (Bissegger et al. 2019; Tobler et al., 2017). Furthermore, complete autosome-to-Y gene duplication will produce M:F coverage = 1.5. Partial Y-gene duplicates will exhibit M:F coverage between 1 and 1.5 (Lin et al. 2022; Tobler et al., 2017; van der Bijl & Mank 2025). To find loci with these characteristics, we first filtered the mapped reads for males and females from the above analysis and called SNPs using SAMTools/bcftools v.19 (Li 2011) *mpileup -Ou -q 20 -Q 20 --skip-indels -a FORMAT/AD,FORMAT/DP* then *call -mv -Oz -f GQ*. SNPs were then filtered using VCFtools v0.1.12b (Danecek et al. 2011) *--maf 0.05 --mac 1 --min-alleles 2 --max-alleles 2 --max-missing 0.9 --min-meanDP 10 --max-meanDP 100 --minGQ 25* and specifying coding regions identified from the Trinity assembled transcriptome.

We used the *--weir-fst-pop* function from VCFtools v0.1.12b (Danecek et al. 2011) to identify the M:F F_ST_ for each SNP in the coding regions. Next, we used the *--site-depth* function to identify the gene read depth for males and females. We used the resulting read depth files to run the *gene_coverage.py* script from Lin et al. (2022) to calculate the M:F coverage of the genes and acquire the corresponding M:F F_ST_. Finally, we calculated M:F SNP density for the putative Y-gene duplications using the sam2pro pipeline (Haubold et al. 2010; http://guanine.evolbio.mpg.de/mlRho/, last accessed September 09, 2024). The M:F read depth of the genes followed a normal distribution, therefore we used two standard deviations to identify genes that have an increased M:F read depth for partial duplicates (>1.206). We bootstrapped the M:F F_ST_ of autosomal genes to calculate the 95% confidence intervals (CI). We called genes that had M:F coverage >1.206 and M:F F_ST_ values greater than the 95% CI (>0.196) as putative Y-gene duplications.

### Gene expression of the putative Y gene duplications

Because expression in the male germ cells is often associated with autosome-to-Y gene duplications (Kaessmann 2010), we assessed expression of putative duplicated genes in *P. reticulata.* Gene-by-cell count matrices from a previously collected single-cell RNA sequencing (scRNA seq) dataset on *P. reticulata* gonads were used to provide expression profiles for male gonad-specific cells (Darolti & Mank 2023). Briefly, gonads were dissected from reproductively mature male guppies and used to generate three nonoverlapping pools with five individuals each. An estimated 8,000 cells from each sample were used to prepare 3’ single-cell libraries using the 10X Genomics Chromium Next GEM Single Cell 3’ kit v3.1. Libraries were sequenced on an Illumina NovaSeq 6000 sequencer, with an average sequencing depth of 20,000 read pairs per cell. CellRanger v5.0.1 (Zheng et al. 2017) was used to build a reference index using the *P. reticulata* genome (Ensembl: GCA_000633615.2; Hunt et al. 2018), align reads and identify cell-associated barcodes to generate gene-by-cell count matrices. Artifactual doublet cells were detected and removed using DoubletFinder v2.0.3 (McGinnis et al. 2019).

Cells were filtered and their counts used for downstream analyses, based on the following quality control thresholds: number of unique molecular identifiers (UMI) per cell (nUMI ≥ 500), number of detected genes per cell (nGene ≥ 250), the log-transformed ratio of genes per UMI (log10GenesPerUMI > 0.80), and the proportion of transcripts mapping to mitochondrial genes (mitoRatio < 0.20). Following quality filtering, cells were clustered into distinct cell types, which were labelled based on known marker gene information (Jiang et al. 2021; Wang et al. 2023).

Genes with low expression - defined as having fewer than one count in at least 10 cells - were removed. Gene counts were summed across cells within each cluster to generate a pseudo-bulk expression matrix. This pseudo-bulk matrix was input into the R package edgeR v4.4.2 (R Development Core Team, 2024) to create a DGEList object. Normalization factors were calculated using the trimmed mean of M-values (TMM) method to scale raw library sizes (*calcNormFactors(DGEList, method = “TMM”*) (Robinson & Oshlack 2010). The resulting dataset contained TMM-normalized counts per million (CPM) values for genes expressed in spermatids and spermatocytes cell types of the *P. reticulata* male gonad. We used these normalized count values to determine cell-specific expression of our putatively Y duplicated genes.

## Results and Discussion

### Genome assembly and transcriptome

Our PacBio sequencing resulted in 41,971,990,442 bp in total, with mean read length of 12.68 Kb, and mean barcode quality of 82 (Supplemental Table 1). Our assembled *de novo* genome was ∼785 Mb (Table 1), comparable to the genomes of *P. reticulata* (731.6Mb; Künstner et al. 2016) and *P. picta* (744 Mb; Metzger et al. 2023). The CANU assembly of the genome had 2,492 contigs with contig coverage ∼ 37X, an NG50 of 3.3 Mb, with contig size ranging up to ∼14 Mb (Table 1). 50% of the assembly resides on 63 contigs (LG50; Table 1).

After orienting the assembly to the *P. picta* female reference genome, we recovered 36 scaffolds with successful alignment to all 23 *P. picta* chromosomes. Our draft genome had a BUSCO completeness of 94.4% and contained ∼ 38% repetitive DNA, as compared to the 29.55% repetitive DNA reported in the *P. picta* reference genome (Metzger et al. 2023; Table 1). The BUSCO score is slightly lower than that reported for *P. picta* (97.65%; Metzger et al. 2023), likely due to the different databases used (cyprinodontiformes_odb10 database in this analysis, eukaryota_odb10 database in Metzger et al. 2023).

We recovered an average of 79.7 million RNA reads per sample (Supplemental Table 1). Following the *de novo* transcriptome assembly, we found a total of 17,521 transcripts and 14,257 of these transcripts BLAST to *P. picta* genome annotation (Table 1). Moving forward, we focused only on the transcripts that had the top BLAST hit to *P. picta* genes.

### Identifying the sex chromosomes and dosage compensation

We recovered an average of 156.0 million Illumina reads per sample, resulting in an average 62.3X coverage (Supplemental Table 1). Using a female (XX) genome, we expect Y degeneration to result in reduced M:F coverage (Darolti et al. 2019), and this approach revealed the *P. bifurca* sex chromosomes to be on guppy chromosome 12. Chromosome 12 had a log_2_ M:F coverage = - 1.0, spanning 0-30Mb, suggesting widespread Y degeneration of this region (Figure 2). There is a small region, 30-34Mb, with equal coverage between males and females (log_2_ M:F ≈0.0), likely representing the pseudoautosomal region (PAR) where recombination still occurs between the X- and Y-chromosome (Figure 2). The significant decrease in male coverage is not observed elsewhere in the genome (Supplemental Figure 1). When compared to the log_2_ M:F coverage of *P. picta* and *P. parae* chromosome 12, we see similar patterns of Y degeneration along the same region of the chromosome (Figure 2).

**Figure 2.**
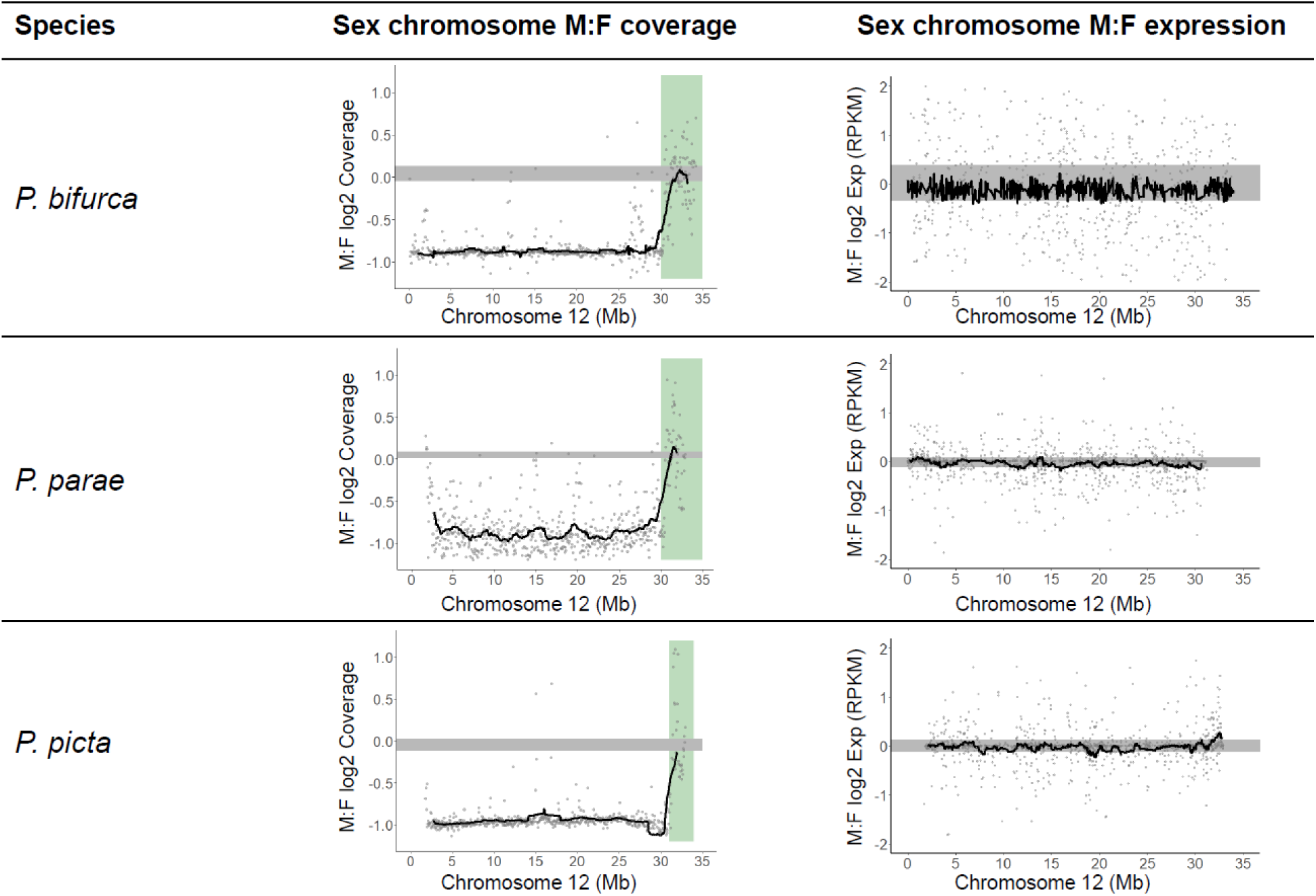
Characterizing the sex chromosome in *P. bifurca* and its closest relatives. Male:female (M:F) log_2_ fold change of reads mapped to chromosome 12, with the value of 0.0 representing equal coverage. The green bar in the middle panel represents the pseudoautosomal region (PAR). The right panel shows the M:F log_2_ fold change of the gene expression of chromosome 12. *P. picta* and *P. parae* reads were mapped to the *P. picta* female reference genome, and *P. bifurca* reads were mapped to the draft *P. bifurca* genome. The black line represents the sliding window with the window size of 50kb. The grey bar represents the 95% CI calculated from 1000 bootstraps of the M:F log_2_ fold change of the autosomes in the respective panels.

We further tested for shared ancestry of the Y chromosome among *P. picta, P. parae* and *P. bifurca* by comparing shared male-specific sequences known as Y-mers (Carvalho & Clark 2013; Kabir et al. 2022; Torres et al. 2018). We found more Y-mers are shared between the three species than expected by chance (measured by the female-specific k-mers), suggesting shared ancestry of the sex chromosome (Supplemental Figure 2).

Although degeneration of Y chromosome gene content leaves males with one functional copy of X-linked genes compared to the two copies present in females, males in *P. picta* and *P. parae* exhibit similar expression of the X in males and females, consistent with complete X chromosome dosage compensation (Darolti et al. 2019; Metzger et al. 2021; Metzger et al. 2023). To test for complete dosage compensation in *P. bifurca*, we compared M:F gene expression on chromosome 12 and found no difference in gene expression between males and females, similar to patterns observed in *P. picta* and *P. parae* and indicative of complete dosage compensation (Figure 2). Complete X chromosome dosage compensation is further corroborated by the allele-specific expression (ASE) distribution, identified by the frequency distribution of the major allele ratio. Female *P. bifurca* show a similar ASE pattern between autosomes and sex chromosomes with no difference in the median major allele ratio (p = 1; Figure 3a), whereas males have a preferential expression from one allele in their sex chromosome (median = 0.767) compared to the autosomes (median = 0.667; p = 0.0276; Figure 3a). Moreover, gene expression between the autosomes and non-recombining region of the sex chromosomes in females (p = 0.493) and males (p = 0.482) do not differ from each other (Figure 3b).

**Figure 3.**
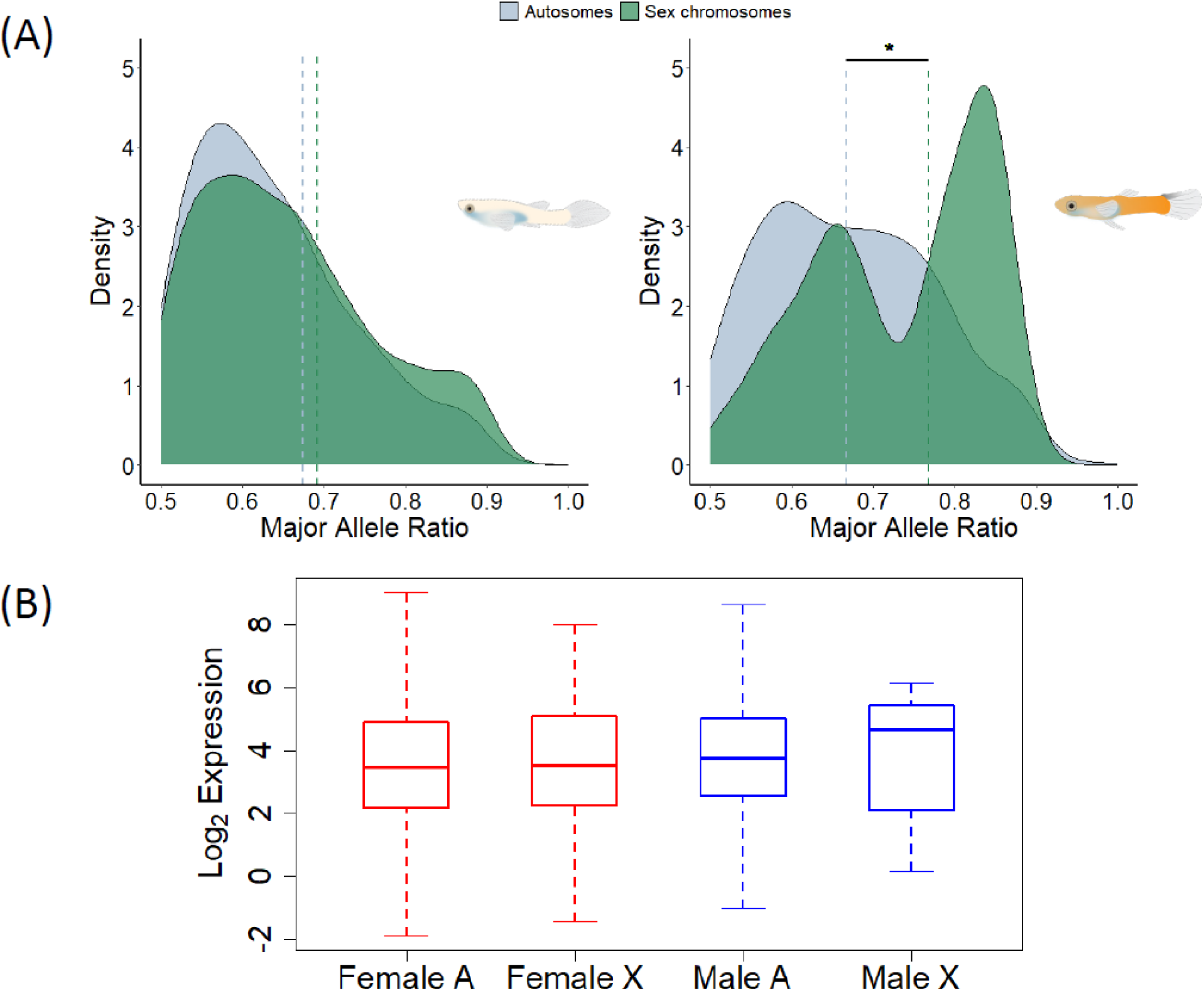
Allele-specific expression (ASE) of *P. bifurca*. A) The density distribution of female (left) and male (right) major allele ratio (frequency) in autosomes (blue) and the sex chromosomes (green). The dashed lines represent the median major allele ratio of each respective distribution. The median value of autosome and sex chromosome major allele ratio in males is statistically significant (Chi-squared test; p = 0.0276). B) Boxplots show the average female (red) and average male (blue) normalized expression between autosomes (A) and the non-recombining region of the sex chromosome genes with an ASE pattern (X) in *P. bifurca*, with no statistically significant difference between the autosomes and X in either sex (Chi-squared test; female p = 0.493; male p = 0.482).

### Putative Y gene duplications

Following Lin et al. (2022), we identified putative autosome-to-Y gene duplications, finding 11 autosomal genes with elevated M:F read depth, elevated M:F F_ST_, and M:F SNP density ≥1, distributed across 8 of the 22 autosomes (Table 2). We used GeneCards (Stelzer et al. 2016) and Gene Ontology Annotation (Ashburner et al. 2000; Carbon et al. 2021) to identify the function of these genes, where known.

**Table 2.** Putative autosome to Y-gene duplications and their autosomal location in *P. bifurca*. Average male:female (M:F) read depth, M:F F_ST_, and M:F SNP density for the coding regions of the gene are reported. Gene functions are referenced from gene ontology (Ashburner et al. 2000; Carbon et al. 2021) and GeneCards (Stelzer et al. 2016). Expression reported is the median of all male samples in CPM if applicable. The gene marked with an * is the top BLAST hit from *P. reticulata* to an uncharacterized gene from the *P. picta* genome annotation.

| Gene Name | Chrom | Mean<br>M:F $F_{ST}$ | M:F Read<br>Depth | M:F SNP<br>Density | Gene Function | Expr. in<br>spermatid<br>(CPM) | Expr. in<br>spermatocyte<br>(CPM) |
| --- | --- | --- | --- | --- | --- | --- | --- |
| *slc4a10* | LG2 | 0.3333 | 1.4561 | 1.0000 | Transporter activity;<br>solute:inorganic anion<br>antiporter activity | N/A | N/A |
| *mapk6* | LG3 | 0.3333 | 1.2185 | 1.0288 | Transferase activity; transferring<br>phosphorus-containing groups;<br>protein tyrosine kinase activity | 62.7700 | 48.3394 |
| *gcshb* * | LG3 | 0.5000 | 1.2079 | 1.1312 | Enzyme binding;<br>aminomethyltransferase activity | N/A | N/A |
| *trub2* | LG5 | 0.5000 | 1.2059 | 1.6863 | Pseudouridine synthase activity | 29.1664 | 15.4314 |
| *ddi2* | LG5 | 0.3182 | 1.2362 | 1.0726 | Aspartic-type endopeptidase<br>activity; regulation of DNA stability; and regulation of<br>protein stability. | N/A | N/A |
| *spsb3* | LG8 | 0.3182 | 1.2727 | 1.0532 | SPRY domain-containing SOCS<br>box protein 3 | N/A | N/A |
| *ccdc97* | LG8 | 0.3333 | 1.4407 | 1.1852 | Coiled-coil domain | 71.5283 | 73.7606 |
| *cfap100* | LG10 | 0.5000 | 1.2326 | 1.4808 | Cilia and flagella associated<br>protein 100 | 21.8748 | 21.0428 |
| *sdc3* | LG11 | 0.5385 | 1.2086 | 1.0000 | Cytoskeletal protein binding | N/A | N/A |
| *ablim3* | LG14 | 0.3333 | 1.2688 | 1.0000 | Actin binding | 8.2903 | 2.1043 |
| *slc7a8* | LG18 | 0.5000 | 1.3852 | 1.5556 | Peptide antigen binding;<br>antiporter activity | N/A | N/A |

We tested the assumption that genes that are expressed in male germ cells are more likely to be duplicated on the Y chromosome and passed on in the male germline (Connallon & Clark 2010; Kaessmann 2010). To do so, we looked at the gene expression of the putative Y-genes in *P. reticulata* spermatocyte and spermatid cells (male germ cell types). We found 5 of the 11 genes had expression in the male gonadal tissues (Table 2). Interestingly, all of the putative Y-gene duplicated genes are expressed in at least one human male testes cell type (Supplemental Table 2).

## Conclusions

In this study, we reveal the conservation of the sex chromosomes in *P. bifurca* and complete X chromosome dosage compensation. Through the use of different sequencing technologies, we built a draft genome with 94.4% completeness and found 14,257 transcript matches to its close relative, *P. picta*. Furthermore, we identified 11 putative autosome-to-Y gene duplications. Our genome is a resource for future research, especially work that uses a comparative approach to better understand the ecology and evolution of *Poecilia* fishes.

## Data availability

All *P. bifurca* sequence data have been deposited under NCBI BioProject PRJNA1259976. The draft *P. bifurca* genome is available under NCBI BioSample SAMN48384351. Data for *P. picta* and *P. parae* used for re-analysis of coverage and expression can be found under NCBI BioProject ID PRJNA528814 and BioProject PRJNA714257, respectively. All scripts and related codes can be found at https://github.com/ljmfong/Poecilia-bifurca-Characterizing-Sex-Chromosome.

## Acknowledgments

This work was funded by a C150 Research Chair and NSERC grants to JEM, a UBC 4^th^ year fellowship to LJMF, and a BRC Informatics Fellowship to BDJ. We would like to thank the Mank Lab for helpful comments and suggestions on the manuscript, especially Y. Lin for genome assembly consultation. We would also like to thank W. van der Bijl for the photo of *P. bifurca*.

---

**Supplemental Table 1.**
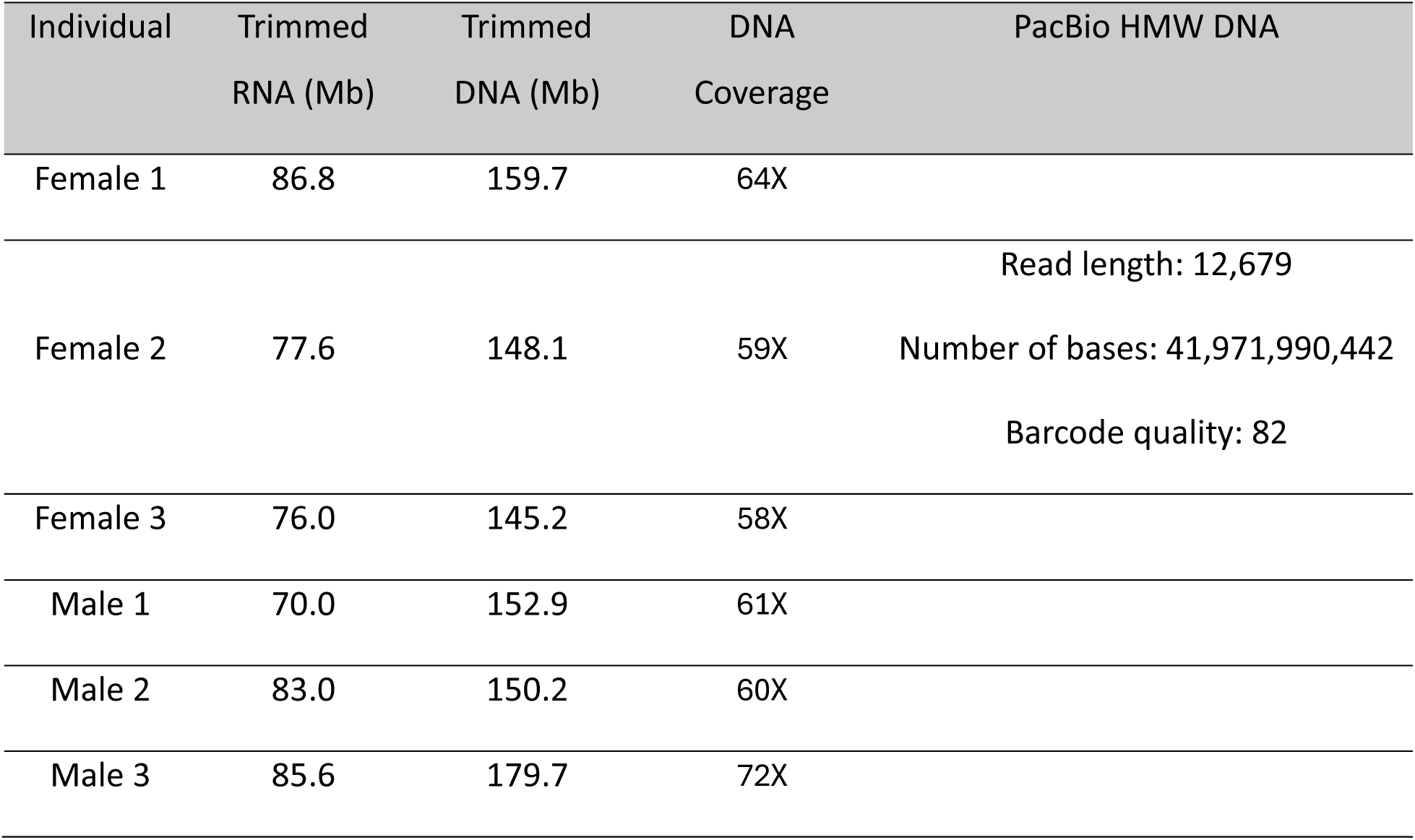
Summary statistics of sequenced *P. bifurca* individuals.

**Supplemental Table 2.**
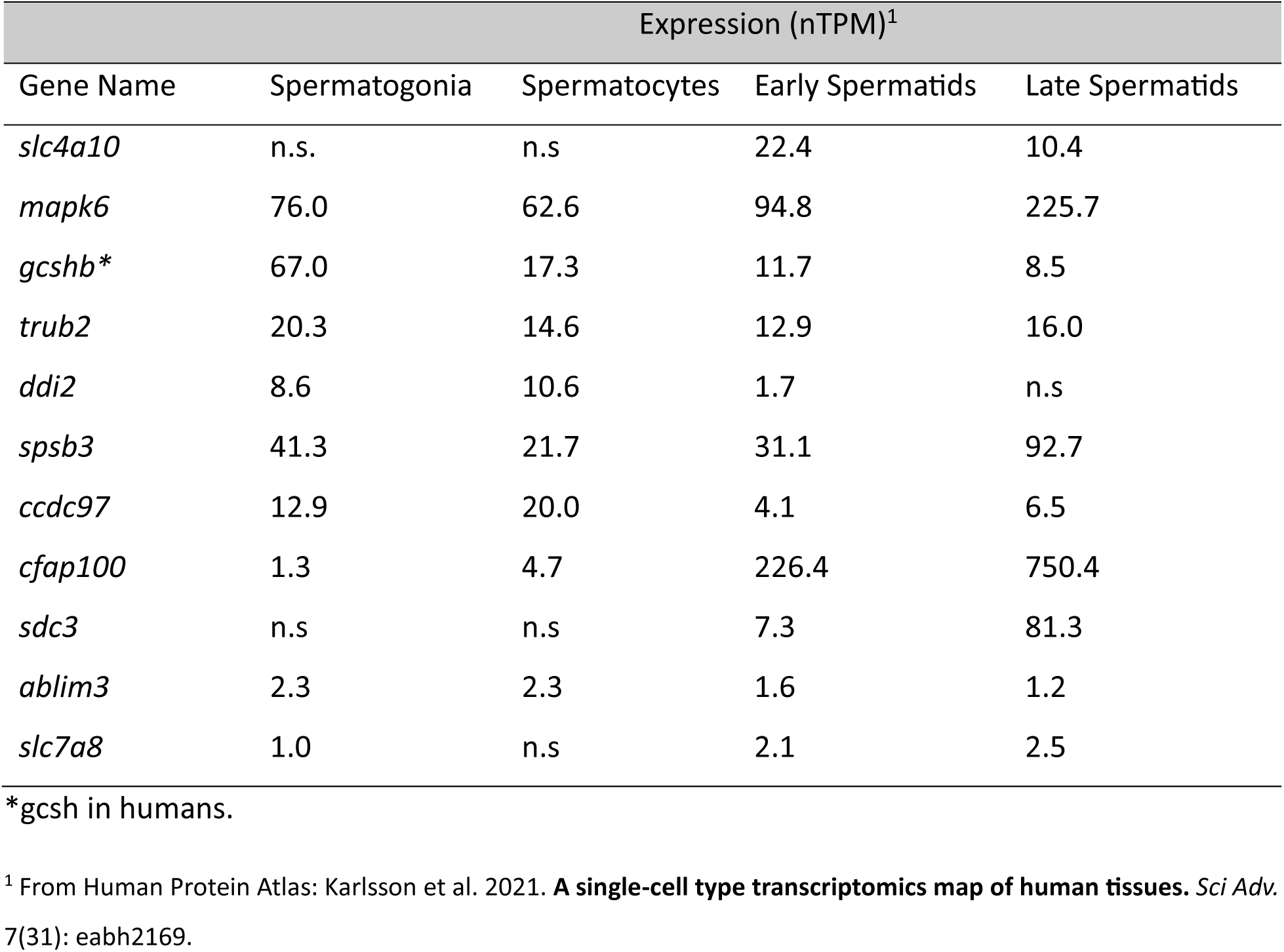
Expression of putative autosome-to-Y duplicated genes in human testes cells.

**Supplemental Figure 1.**
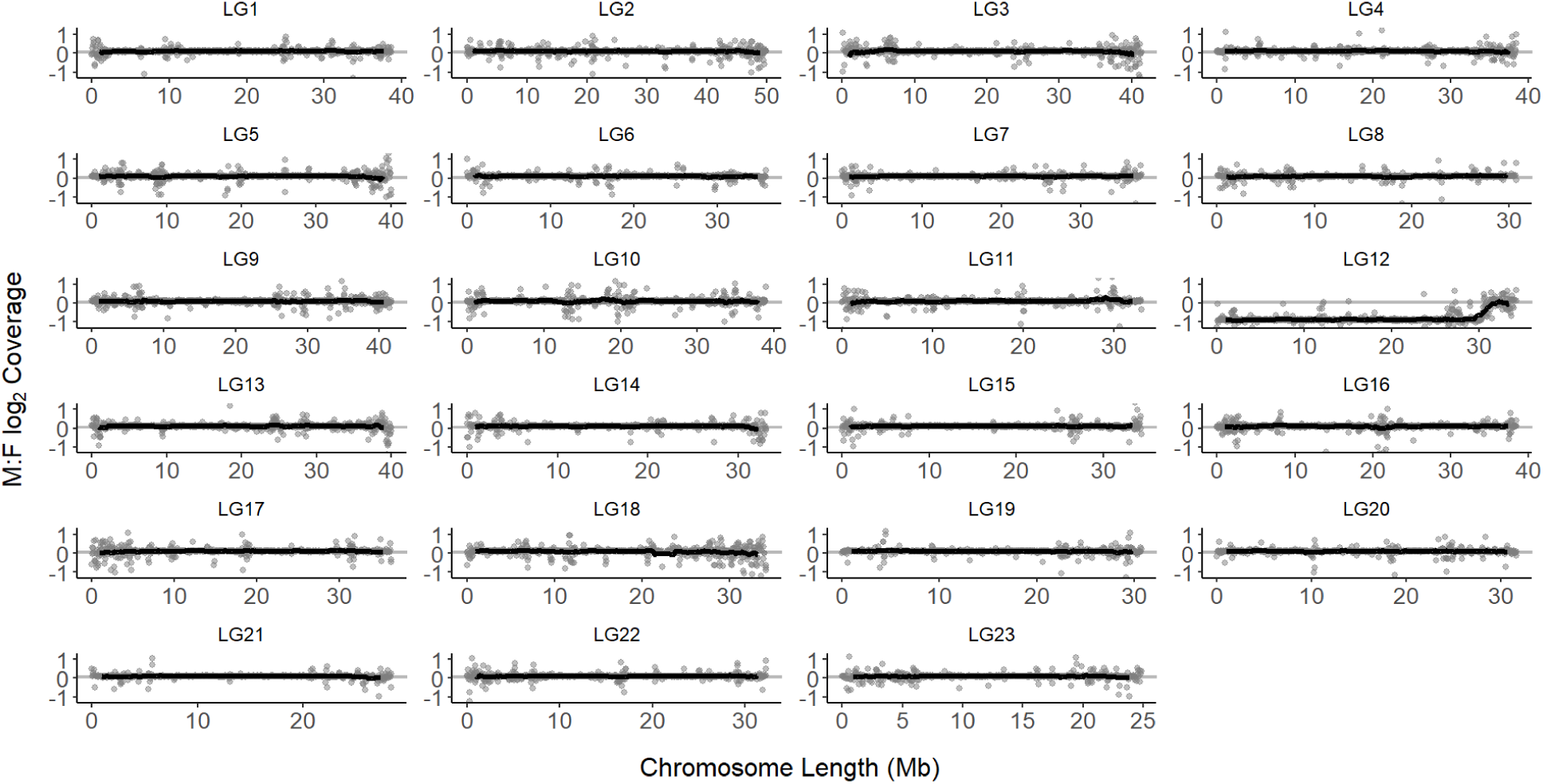
The M:F log_2_ fold change in coverage for all chromosomes in *P. bifurca*. Male and female reads were mapped to the *P. bifurca* draft genome, with the value of 0.0 representing equal coverage. Each panel represents the respective chromosome (LG). The black line represents the sliding window with the window size of 50kb. The grey bar represents the 95% CI calculated from 1000 bootstraps of the M:F log_2_ fold change of the autosomes.

**Supplemental Figure 2.**
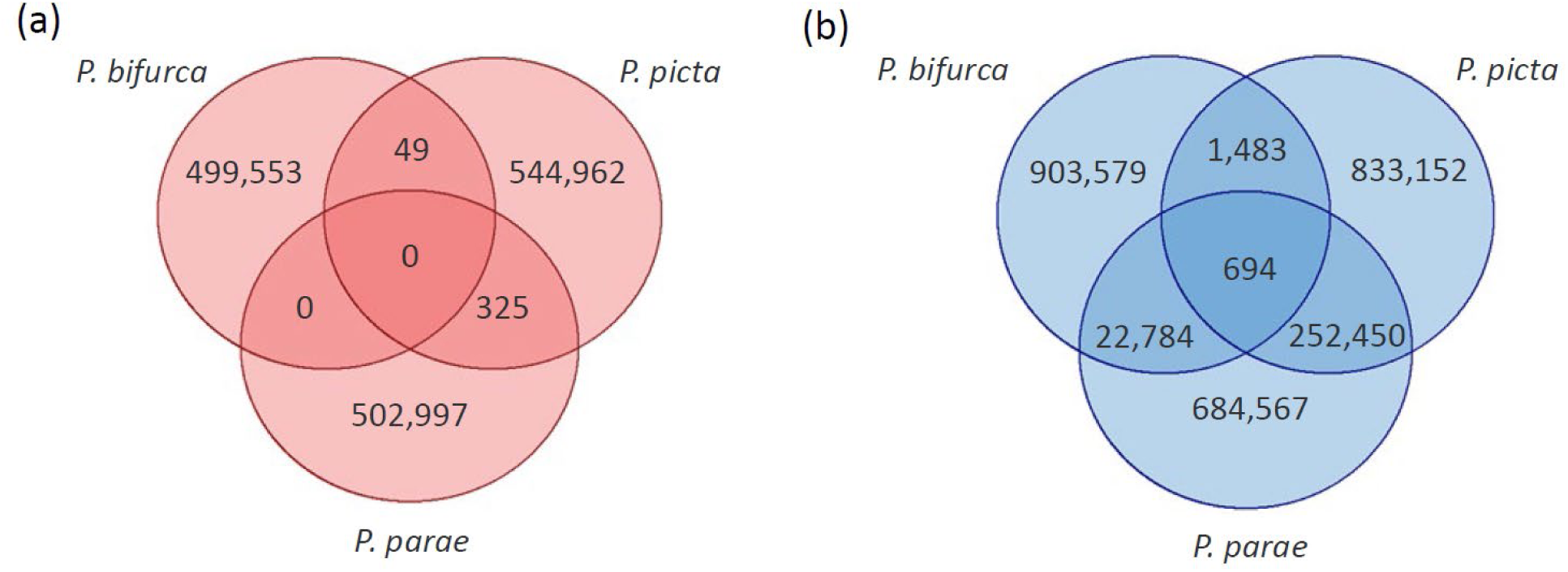
Shared sex-specific k-mers between *P. bifurca*, *P. picta*, and *P. parae*. **a)** Female-specific k-mers found unique in the species and shared between species, representing false positives. **b)** Male-specifc k-mers (Y-mers) found unique in the species and shared between species. There are at least 30X more shared Y-mers than false positives.

